# Sarbecovirus disease susceptibility is conserved across viral and host models

**DOI:** 10.1101/2023.10.11.561544

**Authors:** Sarah R. Leist, Alexandra Schäfer, Ellen L Risemberg, Timothy A. Bell, Pablo Hock, Mark R. Zweigart, Colton L. Linnertz, Darla R. Miller, Ginger D. Shaw, Fernando Pardo Manuel de Villena, Martin T. Ferris, William Valdar, Ralph S. Baric

**Affiliations:** Department of Epidemiology, University of North Carolina at Chapel Hill; Curriculum in Bioinformatics and Computational Biology, University of North Carolina at Chapel Hill; Department of Genetics, University of North Carolina at Chapel Hill; Lineberger Comprehensive Cancer Center, University of North Carolina at Chapel Hill; Department of Microbiology and Immunology, University of North Carolina at Chapel Hill

## Abstract

Coronaviruses have caused three severe epidemics since the start of the 21^st^ century: SARS, MERS and COVID-19. The severity of the ongoing COVID-19 pandemic and increasing likelihood of future coronavirus outbreaks motivates greater understanding of factors leading to severe coronavirus disease. We screened ten strains from the Collaborative Cross mouse genetic reference panel and identified strains CC006/TauUnc (CC006) and CC044/Unc (CC044) as coronavirus-susceptible and resistant, respectively, as indicated by variable weight loss and lung congestion scores four days post-infection. We generated a genetic mapping population of 755 CC006xCC044 F2 mice and exposed the mice to one of three genetically distinct mouse-adapted coronaviruses: clade 1a SARS-CoV MA15 (n=391), clade 1b SARS-CoV-2 MA10 (n=274), and clade 2 HKU3-CoV MA (n=90). Quantitative trait loci (QTL) mapping in SARS-CoV- and SARS-CoV-2-infected F2 mice identified genetic loci associated with disease severity. Specifically, we identified seven loci associated with variation in outcome following infection with either virus, including one, *HrS45*, that is present in both groups. Three of these QTL, including *HrS45*, were also associated with HKU3-CoV MA outcome. *HrS45* overlaps with a QTL previously reported by our lab that is associated with SARS-CoV outcome in CC011xCC074 F2 mice and is also syntenic with a human chromosomal region associated with severe COVID-19 outcomes in humans GWAS. The results reported here provide: (a) additional support for the involvement of this locus in SARS-CoV MA15 infection, (b) the first conclusive evidence that this locus is associated with susceptibility across the *Sarbecovirus* subgenus, and (c) demonstration of the relevance of mouse models in the study of coronavirus disease susceptibility in humans.

---

Over the last three decades, three zoonotic coronaviruses have emerged in humans: SARS-CoV in 2003 (Zhong et al., 2003), MERS-CoV in 2012 (Hijawi et al., 2013) and SARS-CoV-2 in late 2019 (Zhou et al., 2020a, Zhou et al., 2020b). SARS-CoV-2 is the causative agent of the ongoing COVID-19 pandemic. Despite the public health emergency being declared over as of May 11^th^, 2023 (https://www.cdc.gov/coronavirus/2019-ncov/your-health/end-of-phe.html), acute infections with constantly evolving Variants of Concern (VoCs) as well as post-acute sequelae weeks to months after infection pose ongoing global health burdens. Furthermore, it has been shown that coronaviruses in natural animal reservoirs like bats are poised for human emergence (Menachery et al., 2015) and that climate change increases the likelihood of future transmission of viruses from animal to humans (Carlson et al., 2022). Despite a surge in worldwide efforts in understanding disease mechanisms for countermeasure development, many aspects of the disease caused by SARS-CoV-2 specifically, but also sarbecoviruses and other coronaviruses in general, remain elusive. Thus, the need to understand mechanisms driving disease progression, outcome, and ways to intervene remains of high priority.

Murine model systems are one of the most widely used biomedical research tools with which to investigate infectious disease mechanisms and pathogenesis *in vivo*. Mouse-adapted versions of both viruses (SARS-CoV MA15 (Roberts et al., 2007) and SARS-CoV-2 MA10 (Leist et al., 2020) have been used in multiple studies showing their potential to replicate acute respiratory distress syndrome (ARDS) as seen in the human population (Yan et al., 2022, Thieulent et al., 2023, Leist et al., 2020). Even with these available tools, most of these studies focus on one or a few classical mouse strains (e.g. BALB/c or C57BL/6 (Gralinski et al., 2013, Adams et al., 2023)), which neglects the effect of host genetic background on driving disease outcomes. Genetically diverse mouse genetic reference panels allow us to examine host genetic factors contributing to viral disease severity and outcomes.

Host susceptibility loci that regulate coronavirus infection response were first reported with mouse hepatitis virus (MHV) in mice (Ohtsuka and Taguchi, 1997). Over the past decade, we have utilized the Collaborative Cross (CC) genetic reference panel to study a variety of viral pathogens, including the identification of several genetic loci associated with SARS-CoV infectious outcomes (Ferris et al., 2013, Noll et al., 2020, Gralinski et al., 2015, Maurizio et al., 2018, Schafer et al., 2022). The CC is a large panel of recombinant inbred (RI) strains derived from eight genetically diverse founder strains and designed specifically for complex trait analysis and systems genetics approaches (initially described in (Threadgill et al., 2011) and reviewed more fully in (Leist and Baric, 2018)). Herein, we use an F2 cross between coronavirus-susceptible CC006/TauUnc and coronavirus-resistant CC044/Unc (hereafter, CC006 and CC044, respectively) mice to more thoroughly investigate the genetic architecture underlying both SARS-CoV and SARS-CoV-2 disease outcomes, as well as assess concordance with a pre-emergent (HKU3-CoV) virus.

We purchased a set of 10 CC strains (listed in Fig 1.A-C) from the System’s Genetics Core Facility (SGCF) at UNC, and screened them for divergent responses to SARS-CoV MA15 infection. Four 10-week-old female mice from each of these ten CC strains were intra-nasally infected with 10^4^ plaque-forming units (PFU) of mouse-adapted SARS-CoV MA15 (Roberts et al., 2007), and changes in body weight were monitored for four days post-infection (dpi). Lung congestion scores as a gross evaluation of histopathological changes at the time point of harvest as well as viral titer in lungs on day 4 were measured. Viral titers were determined by plaque assay, as previously described (Schafer et al., 2022). We observed highly variant disease outcomes in terms of weight loss (**Fig. 1A**), congestion scores (**Fig. 1B**) and viral titer (**Fig. 1C**) across these ten strains. From this screen we identified CC006 as susceptible and CC044 as resistant to severe disease. CC006 exhibited a severe disease response marked by 50% mortality and 19% loss of body weight on average in surviving mice by day 4 post-infection. In contrast, CC044 mice were resistant to disease, with no mortality and little if any change in body weight on average. The overall weight loss trajectories (area above the curve, AAC) of CC006 and CC044 mice were significantly different from each other (p = 6×10^−3^, Tukey’s HSD). CC006 mice also exhibited significantly higher congestion scores than CC044 mice (p = 1×10^−4^, Tukey’s HSD test). Importantly, while these strains had divergent disease responses, they showed similar viral loads (p = 0.99, Tukey’s HSD test), suggesting that variation in disease severity is due to variable host response rather than variation in viral burden. Following the initial screen, we repeated this experiment with CC006, CC044, and CC023/GeniUnc and validated their divergent responses (**Fig. 1D-F**). Weight loss trajectories (**Fig. 1D**, p = 3×10^−4^) but not congestion scores (**Fig. 1E**, p = 0.13) were significantly different from each other between CC006 and CC044 in this repeat experiment. As in the original screen, titer was not significantly different between CC006 and CC044 mice (**Fig. 1F**, p = 0.53). As such, CC006 and CC044 seemed strong candidates to generate a targeted F2 cross for genetic mapping.

**Figure 1.**
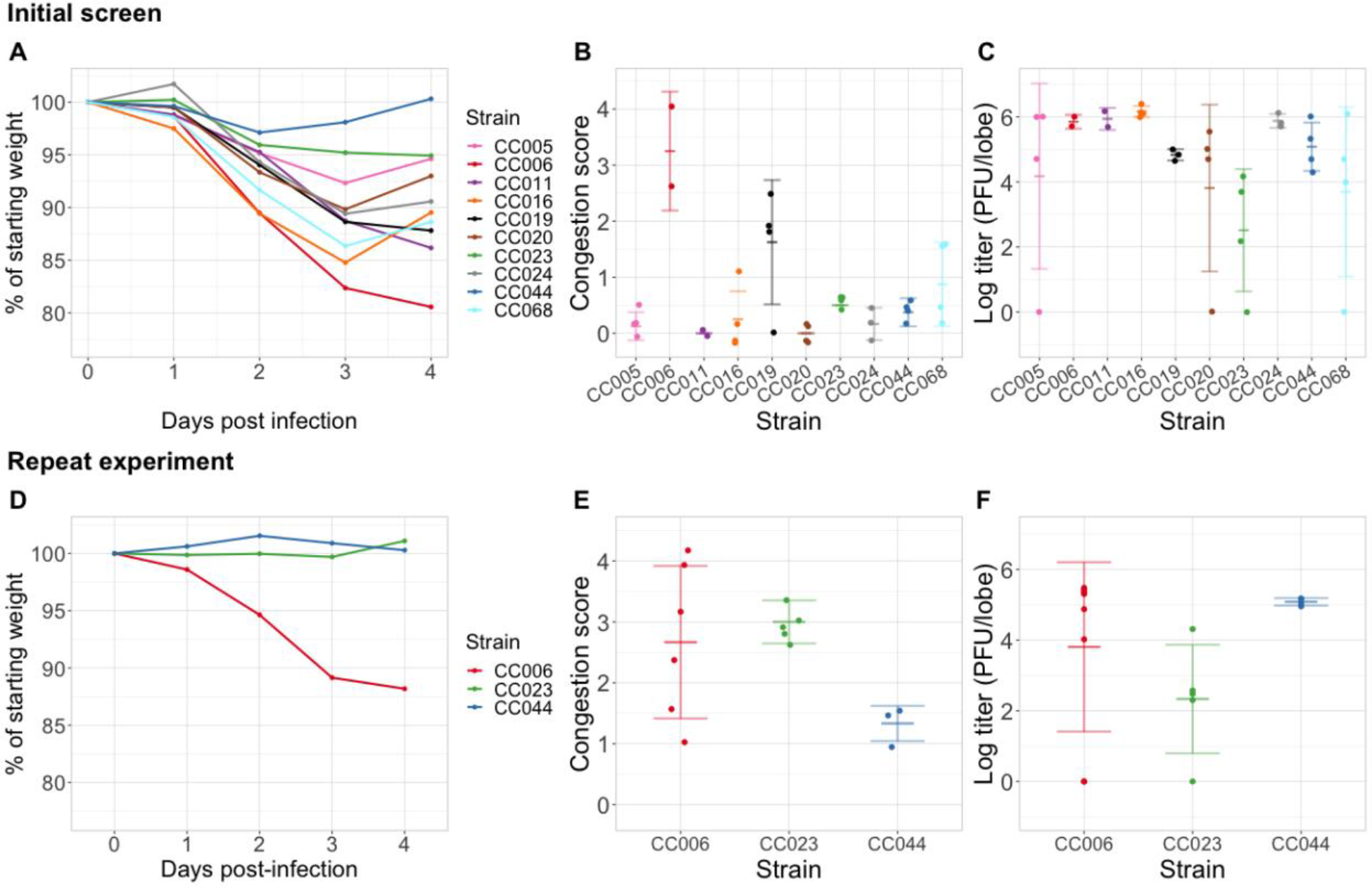
CC strains demonstrate variable response to SARS-CoV MA15 infection. A) Weight loss for four dpi, B) lung congestion score, and C) log-transformed viral titer following infection with SARS-CoV MA15 in four female mice from each of 10 CC strains. D) Weight loss, E) lung congestion scores and F) log-transformed viral titer in four female mice from each of three CC strains, chosen for their divergent responses in the initial screen.

We generated a genetic mapping population of CC006xCC044 F2 mice, using a balanced breeding design as we have previously described (Schafer et al., 2022, Gralinski et al., 2017). In total, we generated 755 F2 mice from all four grandparental combinations. These F2 mice were exposed to one of three coronaviruses: the clade Ia sarbecovirus SARS-CoV MA15 (n=391), the clade Ib sarbecovirus SARS-CoV-2 MA10 (n=274) and also the clade II sarbecovirus HKU3-CoV (HKU3-CoV MA) (n=90), a mouse-adapted bat virus (Becker et al., 2008). We assessed disease severity and progression by measuring weight loss for four days following infection, and at 4 dpi, we euthanized the mice and assessed their overall lung pathology in the form of a congestion score. Weight loss following SARS-CoV MA15 (**Fig. 2A**) and SARS-CoV-2 MA10 (**Fig. 2B**) infection in these F2 mice expanded the range of what was observed in the parental strains following SARS-CoV MA15 infection (**Fig. 1A**), as is often observed in our mapping crosses (Schafer et al., 2022, Smith et al., 2016, Gralinski et al., 2017) (). F2 mice infected with SARS-CoV MA15 lost an average of 5% (ranging from a 7% gain to 26% loss) of their body weight, with congestion scores between 0-4 (median = 0.5). F2 mice infected with SARS-CoV-2 MA10 lost an average of 6% (ranging from 33% gain to 28% loss) of their body weight, with congestion scores between 0-4 (median = 0.5). F2 mice infected with HKU3-CoV MA exhibited less severe weight loss and congestion scores compared to those infected with SARS-CoV MA15 and SARS-CoV-2 MA10. There was still a substantial range of disease responses after HKU3-CoV MA infection, with mice losing an average of 1% (ranging from 7% gain to 13% loss) of their body weight by day 4 and congestion scores ranging from 0-2 (median = 0.5). These results highlight the value of our mouse-adapted viruses, SARS-CoV MA15 and SARS-CoV-2 MA10, in leading to highly correlated parameters of disease manifestation. The evaluation of coronavirus-induced disease across three different viruses indicates that there might be host genetic mechanisms that lead to common disease responses despite genetic differences across coronaviruses, and thus increases the impact of the findings to potentially encompass future viral epidemics.

**Figure 2.**
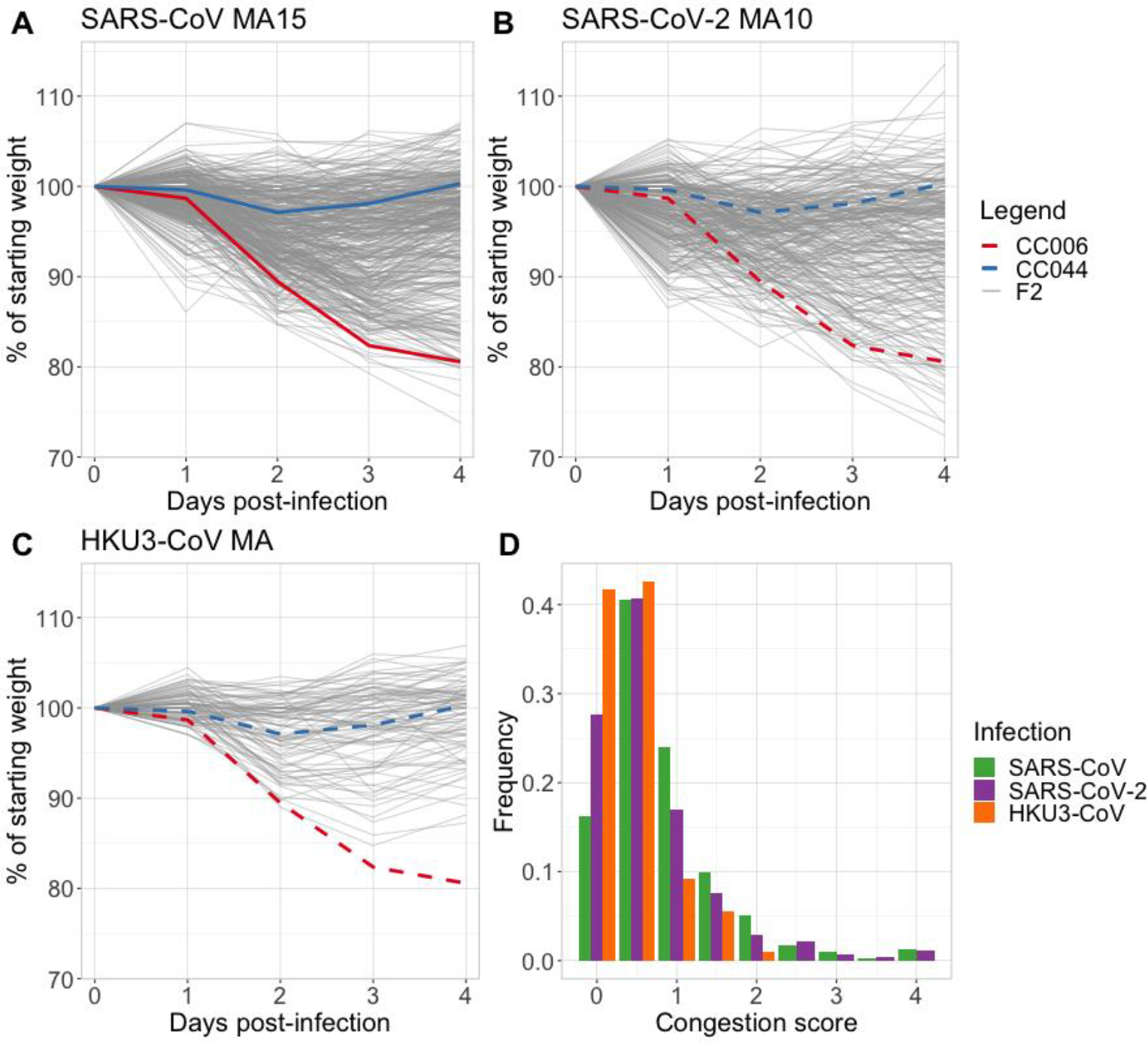
Disease phenotypes after SARS-CoV MA15, SARS-CoV-2 MA10, and HKU3-CoV MA in CC006xCC044 F2 mice. (A) Weight loss in n=391 F2 mice after infection with SARS-CoV MA15. (B) Weight loss in n=274 F2 mice after infection with SARS-CoV-2 MA10. (C) Weight loss in n=90 F2 mice after infection with HKU3-CoV MA. Red and blue lines in A-C represent average body weight trajectory of CC006 and CC044 mice, respectively, after infection with SARS-CoV MA15 (see Fig. 1A). (D) Distribution of congestion scores in F2 mice, colored by infection group.

Concurrent with viral challenge, we genotyped these F2 mice on the miniMUGA array (Sigmon et al., 2020). We filtered the 10,819 biallelic single nucleotide polymorphisms (SNPs) down to a set of 2,599 well-performing markers which segregated between CC006 and CC044 on the autosomes and X-chromosome. These markers were well-distributed across the genome, with a median and maximum distance between markers of 0.5 Mb and 50 Mb, respectively. Using this set of markers, we performed QTL mapping using R/qtl (Broman et al., 2003) in both the SARS-CoV MA15- and SARS-CoV-2 MA10-infected mice. For each virus, mapping was performed on five phenotypes: weight on each of days 2-4 (with weight on day 0 as a covariate to control for baseline weight), an “area above the curve” (AAC) measure to capture the overall weight loss trajectory for each mouse, and congestion score. We identified a total of seven loci associated with these aspects of coronavirus disease severity (**Table 1**), six significant loci (*HrS43-48*, P < 0.05) and 1 suggestive locus (*HrS49*, P < 0.10).

**Table 1.**
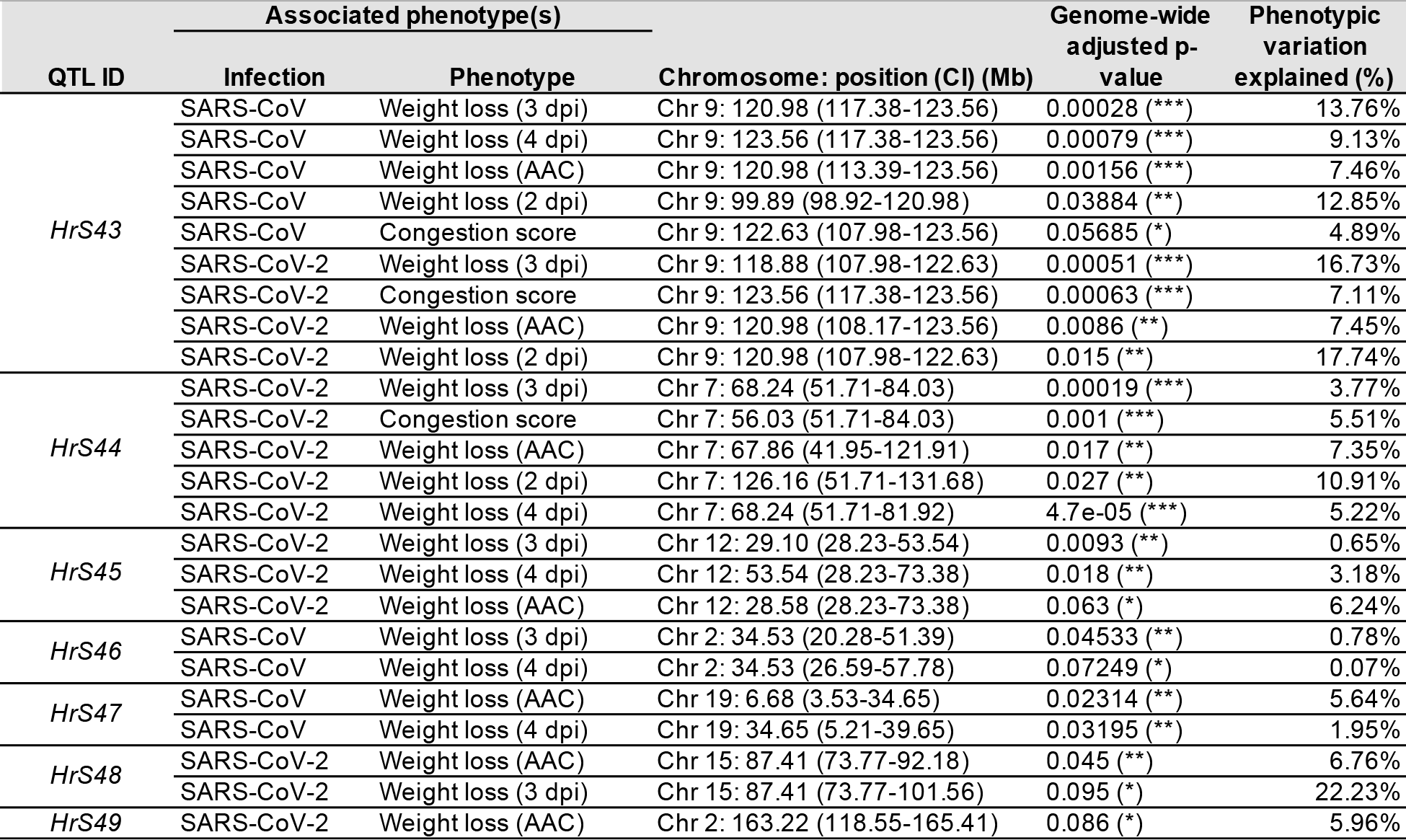
Quantitative trait loci identified in CC006xCC044 F2 mice. Next to genome-wide adjusted p-values, *** indicates highly significant (P < 0.005), ** indicates significant (P < 0.05) and * indicates suggestive (P < 0.10).

*HrS43* overlaps with a locus we had previously identified, *HrS26*, that was associated with multiple disease traits following SARS-CoV MA15 infection in a CC011xCC074 F2 cross (Schafer et al., 2022). As with *HrS26, HrS43* is associated with both weight loss and congestion scores following SARS-CoV MA15 infection (**Fig. 3A-B, Fig. 4A**). We also show that *HrS43* is associated with weight loss and congestion scores following SARS-CoV-2 MA10 (**Fig. 3C-D, Fig. 4B**) infection. Mapping this locus in two independent studies strengthens support for this locus as a driver of disease. Although we lacked sufficient sample size for QTL mapping in the HKU3-CoV-MA-infected mice, we asked whether there was evidence that *HrS43* also impacts HKU3-CoV MA infection outcome. *HrS43* was associated with both weight loss on day 2 (P=0.01) and weight AAC (P=0.05, **Fig. 4C**). Other QTL associated with disease outcomes following HKU3-CoV MA infection were *HrS44* (weight loss on day 3 (p=0.02), day 4 (p=0.05), and AAC (p=0.02)) and *HrS46* (congestion score (p=0.03)).

**Figure 3.**
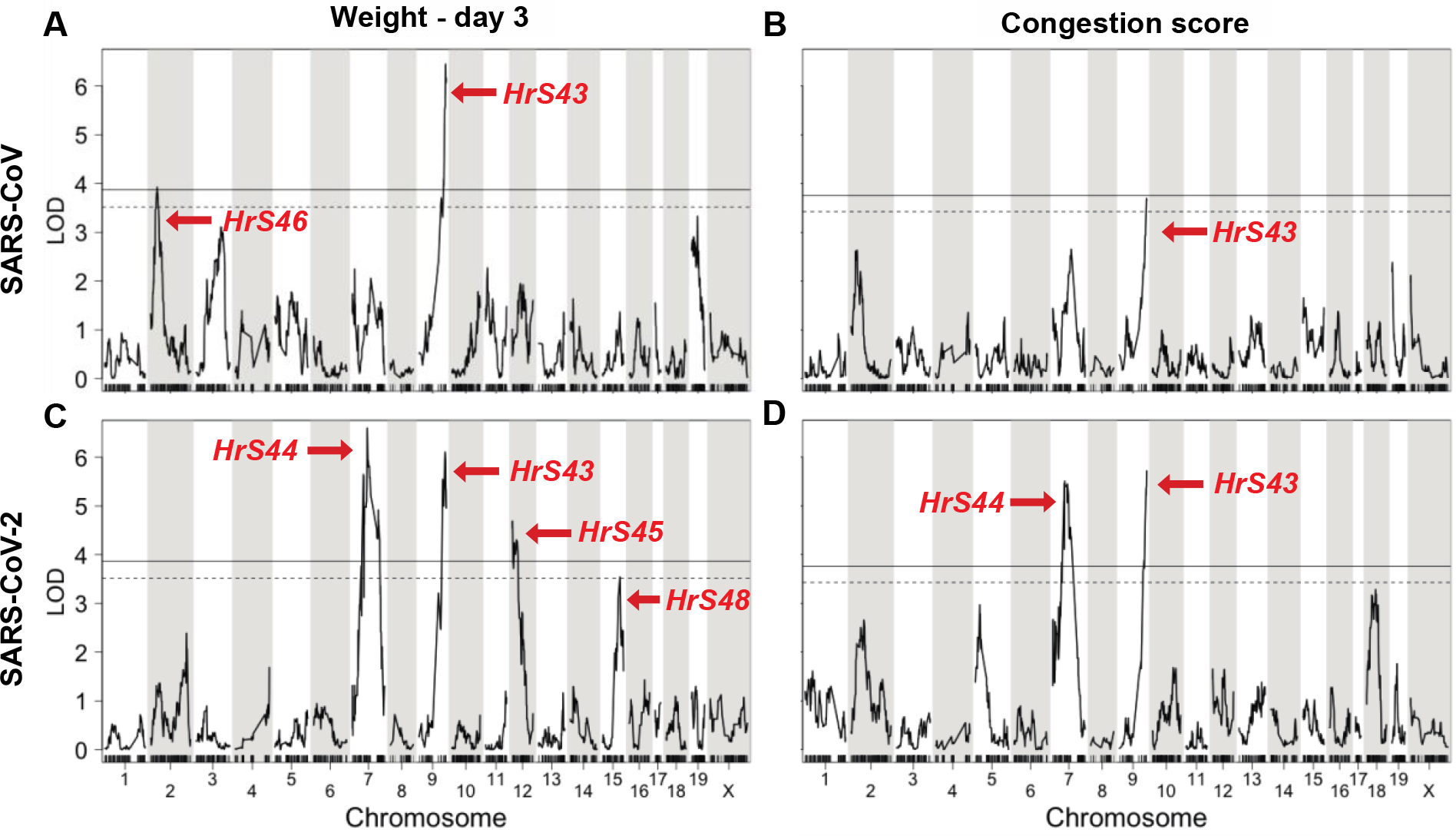
Genome scans from QTL mapping on A) weight loss on day 3 and B) congestion score in SARS-CoV MA15-infected F2 mice, and C) weight loss on day 3 and D) congestion score in SARS-CoV-2 MA10-infected F2 mice. Solid and dashed lines represent 5% and 10% significance thresholds, respectively. *HrS43* overlaps with *HrS26* previously mapped for SARS-CoV MA15 in CC011xCC074 F2 cross.

**Figure 4.**
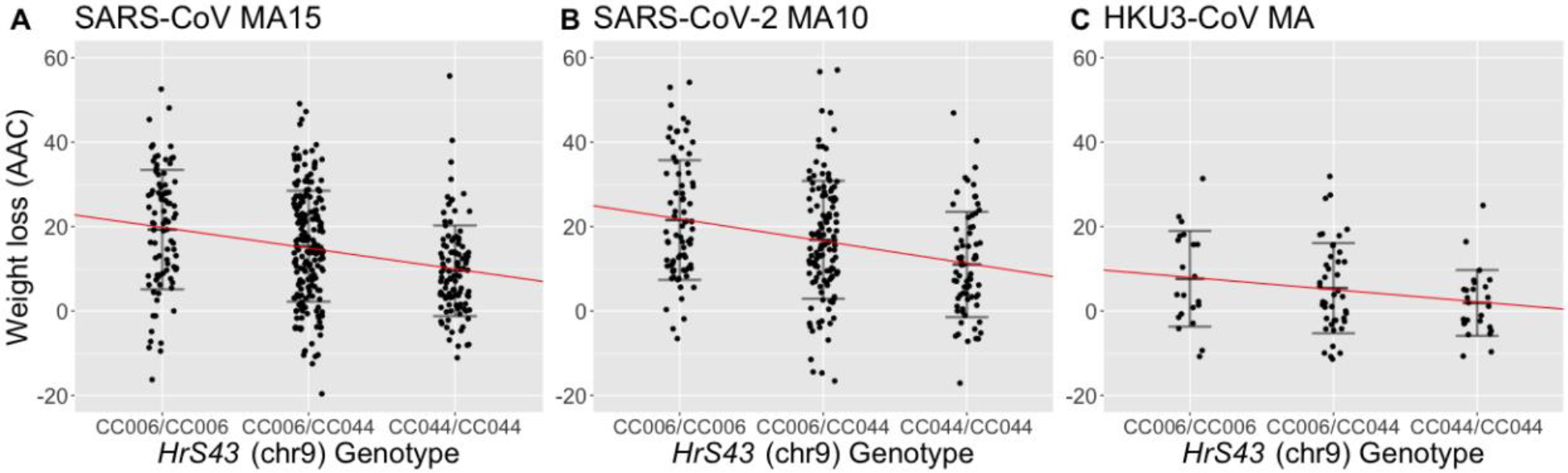
Relationship between genotype at *HrS43* peak marker S3N094839317 and weight loss (AAC) in A) SARS-CoV MA15-infected mice (genome-wide adjusted P = 1.6×10^−3^), B) SARS-CoV-2 MA10-infected mice (genome-wide adjusted P = 8.6×10^−2^), and C) HKU3-CoV MA-infected mice (P = 0.05).

Consistent with other studies (Schafer et al., 2022, Gralinski et al., 2017, Gralinski et al., 2015), we have shown that response to sarbecovirus infection within this cross is influenced by multiple loci. Importantly, our results here provide evidence, first suggested in our prior study (Schafer et al., 2022), that *HrS26*/*HrS43* affect susceptibility via a pan-coronavirus mechanism. Importantly, this locus also appears to be evolutionarily conserved across mammals, as the syntenic region in humans was associated with severe COVID-19 disease in several GWAS (Covid-19 Host Genetics Initiative, 2021, Severe Covid-19 GWAS Group, 2020, Pairo-Castineira et al., 2021, Shelton et al., 2021). Given the overlap between *HrS26* and *HrS43*, and the apparent cross-species and cross-coronavirus relevance, we sought to use genetic data from both crosses to improve our assessment of candidate genes. In our previous study (Schafer et al., 2022), we reported genes that had variants segregating between the strains involved in that study, CC011 and CC074. Here, since the founder strain haplotypes segregating between CC011 and CC074 are different than those between CC006 and CC044, we can provide further refinement by looking for candidate genes with variants segregating between the strains in both crosses. We categorize candidate genes as those with shared SNPs that indicated a common variant affecting susceptibility and those with SNPs that are not shared but point to common susceptibility genes. This approach allowed us to refine potential candidate genes from 971 within the locus to 304. We also report genes that are indicated in one cross but not the other (i.e. variants suggesting genetic pleiotropy).

Using this refined analysis, we cross-indexed our results with several genes that have been associated with severe COVID-19 in human GWAS (Coperchini et al., 2020). These studies have pointed to a cluster of six genes located on human chromosome 3 (syntenic with distal mouse chromosome 9, where *HrS26* and *HrS43* localize): *SLC6A20, LZTFL1, CCR9, FYCO1, CXCR6*, and *XCR1*. In our earlier study, we demonstrated that at least two of the genes underneath this locus (*Cxcr6* and *Ccr9*) contributed to these disease responses. Besides the six genes associated with severe COVID-19, we identified variants of several other genes within *HrS43* with involvement in immunological and antiviral processes and which have been shown to specifically have a role during COVID-19 development and disease. Several chemokine receptors, like *Ccr4, Ccr8, and Cx3cr1*, are involved in shaping the specific immune (adaptive) responses during SARS-CoV-2 infection (Wang et al., 2022, Zhang et al., 2022, Zhou et al., 2021, Victor et al., 2022), while allelic variants (*Cmtm8, Cspg5, Dbr1, Gnai2*, and *Mlh1*) have been described to have functions in DNA splicing, stability, and maintenance during COVID-19 disease (Chen et al., 2021, Ariumi, 2022). Interestingly, we also identified two genes, *Dcaf1*, a regulator of TMPRSS2 expression, and *Xrn1*, an exonuclease, two cellular genes, which are directly involved in the SARS-CoV-2 replication (Chen et al., 2021, Ariumi, 2022).

Here, we show that several disease phenotypes during SARS-CoV MA15, SARS-CoV-2 MA10 and HKU3-CoV MA infection in a CC006xCC044 F2 cross are associated with a locus (*HrS43*) which is also associated with SARS-CoV-2 disease outcome in humans. As such, this study is the first to conclusively demonstrate that sarbecovirus disease is, in part, controlled by common host loci, and that this control is relevant across mammalian hosts. Such conserved and broadly relevant loci are of high interest for global public health outcomes, as they point to mechanisms of susceptibility that can inform treatment and intervention strategies for current and future viral epidemics. Further, consistent associations across mouse and human studies demonstrate that susceptibility mechanisms are conserved across mammals and supports the use of mouse models in the study of host genetic response to viral infection. Altogether, our studies point to a set of host genes driving these outcomes across both human and mouse systems, suggest that future virus emergences can be putatively treated based on prior emergence analyses, and reinforce the utility of mammalian model systems to inform our understanding of emerging infections.

---

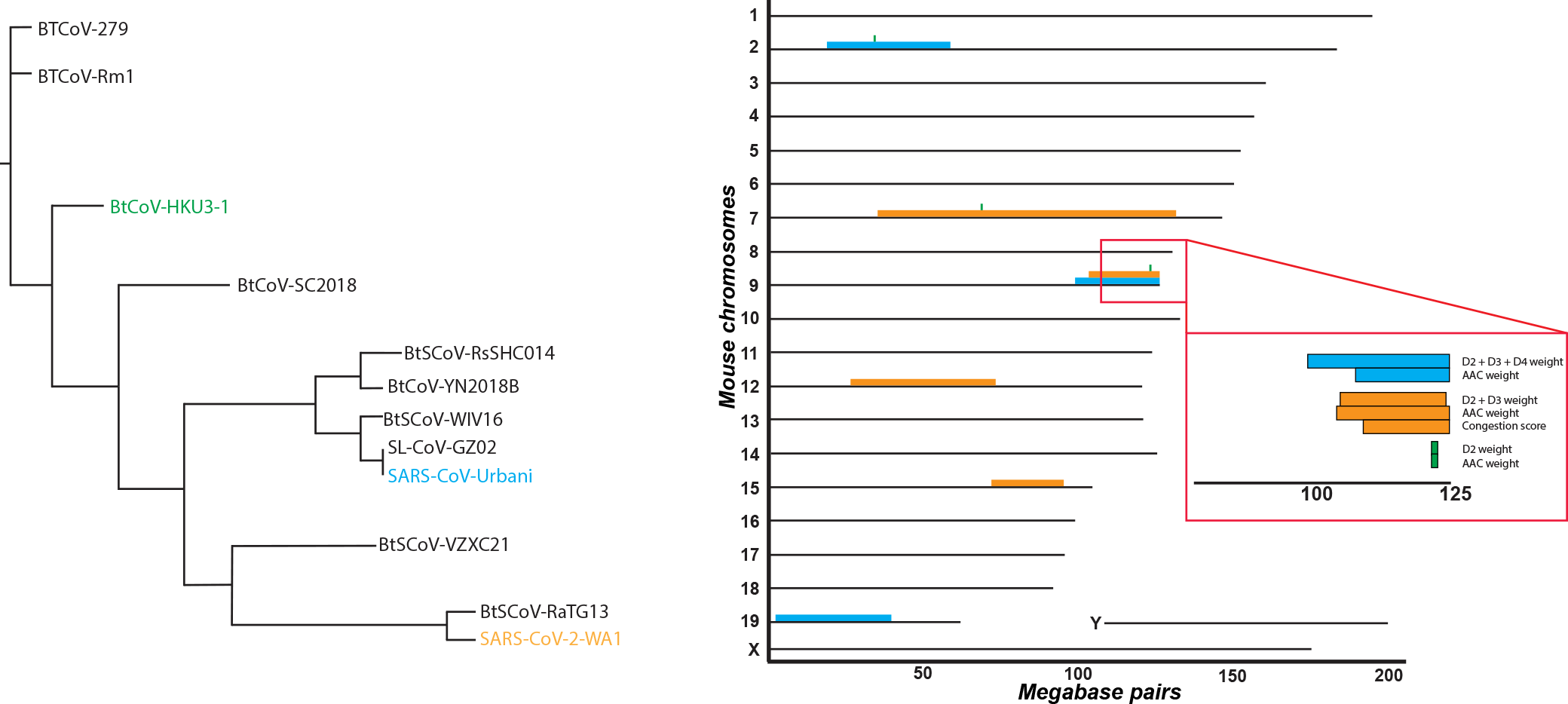

